# Climatic niche dynamics and its role in the insular endemism of *Anolis* lizards

**DOI:** 10.1101/208652

**Authors:** Julian A. Velasco, Enrique Martinez-Meyer, Oscar Flores-Villela

## Abstract

We evaluated the tempo and mode of climatic niche evolution in the radiation of Caribbean *Anolis* lizards and the role of climate in shaping their exceptional insular endemism. Using phylogenetic comparative methods, climatic niche data and a calibrated phylogeny, we reconstructed climatic niche dynamics across time and space for Caribbean *Anolis* lizards. We found evidence of several instances of niche shifts through the Caribbean *Anolis* radiation. Caribbean anole species have diversified mainly along a precipitation rather than a temperature gradient. Only a few lineages have colonized both cold and hot conditions. Furthermore, most of the single-island endemic species are climatically restricted to its native islands and a small set of species might the potential to colonize other islands given its climatic niche requirements. Overall, we found evidence that climate niche conservation has played a role structuring current insular *Anolis* endemism. The observed climatic dissimilarity across the Greater Antilles likely limit successful population establishment of potential exotic insular species.

## Introduction

Insular systems are a well-known example of unique diversity in biogeography, one that is reflected in an elevated number of endemic species (MacArthur & Wilson, 1967; Losos & Ricklefs 2010). High insular endemism results from a combination of factors, such as colonization, speciation, island size, isolation and environmental uniqueness (Kier *et al.*, 2009). Recently, some studies started to take speciation into account as the main driver of island endemism (Kisel & Barraclough, 2010; Qian & Ricklefs, 2012). However, little is known about the role of ecological niche dynamics, understood as the balance of niche conservatism (i.e., the tendency of species to retain ancestral ecological niche traits; Wiens & Graham, 2005) and niche shifts on shaping patterns of island diversity and endemism (Hosner *et al.*, 2014).

Ecological niche theory predicts that species’ geographical ranges at coarse-grain scales are constrained by a set of abiotic and biotic factors, and dispersal limitations (Peterson *et al.*, 2011). For insular species, geographical range is thought to be constrained by strong dispersal barriers rather than biotic or environmental factors (Brown & Lomolino, 1998). However, few tests have been conducted to disentangle the importance of geographical and ecological barriers on the range limits of insular endemic species. Consequently, it is still unknown whether single-island endemic species might encounter suitable ecological niche conditions outside their native areas and therefore what factors drive their current distribution in insular systems.

Niche conservatism and niche shifts patterns emerge as two non-mutually exclusive patterns from ecological niche dynamics along time and across space (Donoghue & Edwards, 2014). These two patterns might contribute to explain the high degree of endemism in many insular systems. For instance, if insular diversity is shaped only by dispersal events, the species colonizing an island might exhibit similar climate requirements (i.e., an environmental filter can limit dispersal events). This niche similarity might either be the result of niche conservatism or niche convergence because of evolving towards similar climate regimes (Boucher *et al.*, 2011). In contrast, if insular diversity is shaped only by *in situ* speciation, species might exhibit higher niche similarity because of niche conservatism and niche shifts might eventually occur because of climatic opportunity (Algar & Mahler, 2015). In addition, for insular clades some recent studies have found evidence of a direct link between climate and diversification (Algar & Mahler, 2015; Velasco *et al.*, 2016), but still it is not clear how phylogenetic niche conservatism (PNC) or niche divergence pattern can mediate species diversification and ultimately shape species richness and endemism patterns. Using modern phylogenetic comparative methods, it is possible to discern between the two evolutionary scenarios: niche conservatism and niche shifts (Mahler *et al.*, 2013; Bravo *et al.*, 2014). Although there is theoretical evidence that multiple evolutionary processes can drive niche similarity (Revell *et al.*, 2008), the distinction between these two patterns is a crucial step toward understanding how ecological niches evolved at coarse scales and how the two patterns affected insular species richness and endemism patterns.

*Anolis* lizards offer a well-known example of adaptive radiation in both insular and mainland settings (Losos, 2009). Insular anole diversity in the Caribbean basin is explained by a combination of geographic, ecological and evolutionary factors (Losos, 2009). Caribbean anole species are notably endemic to a single island with a few species having colonized other islands mediated by human activities (Helmus *et al.*, 2014). This exceptional endemism is reflected in high species turnover rates between islands and is explained by geographic and environmental factors (Stuart *et al.*, 2012). However, climatic niche dynamics might help explain how these exceptional patterns emerged. While it is known that dispersal and available climate space can drive species diversification in insular systems (Bacon *et al.*, 2012; Wuest *et al.*, 2015; Velasco *et al.*, 2016), these factors have not been explored as drivers of *Anolis* endemism in the Greater Antilles.

Here, we evaluate how climatic niches have evolved through time and space in Caribbean *Anolis* lizards and how niche continuum process (niche conservatism-niche divergence) can help to explain the exceptional endemism observed today. We address whether anole lineages have converged to similar climate regimes in different islands (niche convergence) and how niche disparity accumulated through radiation. In addition, we evaluated the degree to which climate has played a role in shaping current distributions of single-island endemic species.

## Materials and Methods

### Phylogenetic estimation and divergence times

We used a time-calibrated (MCC-tree) phylogeny from Poe *et al.*, (2017). We excluded all mainland species from this tree and the pruned Caribbean tree was composed of 141 species from a total of 166 currently recognized for the Caribbean islands (Poe *et al.*, 2017). The remaining 25 species were not included in this tree due to the lack of molecular data. Anole trees are available from http://www.stevenpoe.net/publications.html.

### Climatic niche convergence and niche disparity through time

We compiled the occurrence records for 141 Caribbean *Anolis* from an updated database of 24,074 locality records for *Anolis* species compiled from several sources (e.g., Algar & Mahler, 2015; Global Biodiversity Information Facility -GBIF-; and original descriptions). We careful revised each species and eliminated erroneous and doubtful records. We extracted climate information for annual mean temperature (bio1) and annual precipitation (bio12) from the WorldClim database (Hijmans *et al.*, 2005) for the locality record of each species and calculated the average for species. We used only these two variables to build the occupied climate space for Caribbean *Anolis* lizards. We preferred to use these two direct variables instead scores from ordination methods (e.g., Principal Component Analyses -PCA-) because the interpretation of occupied climate space is straightforward.

We tested whether climatic niche variation in Caribbean anole lineages is consistent with a scenario of adaptive convergence. We used the SURFACE program (Ingram & Mahler, 2013), which uses a phylogeny and niche traits, to fit an Ornstein-Uhlenbeck (OU) model in which lineages evolved toward convergent climate regimes (adaptive peaks) in a macroevolutionary landscape. An OU model describes an evolutionary process that includes two components: a deterministic tendency to evolve toward optimal states (i.e., adaptive regimes), and a stochastic component that is interpreted as evolutionary changes by natural selection and genetic drift (Butler & King, 2004). The method makes it possible to identify the maximum number of convergent adaptive peaks without any *a priori* delimitation. SURFACE starts with a model with a single adaptive regime for all species and then increases the number of adaptive regimes using a stepwise model selection procedure based on the corrected Akaike Information Criterion (AICc). A new peak shift is added at each step to improve the fit of the model (forward phase). In the backward phase, the final set of previously identified regimes is collapsed to find further improvements of model fit. SURFACE finds the maximum number of adaptive regimes that best fit the data and identifies which clades are convergent towards the same adaptive peaks. We then tested whether our convergent pattern for climatic niche differed from one expected by a null Brownian motion model (BM). We compared our observed convergence parameter (delta k) with 500 simulated delta k under a BM process. We also evaluated the effect of phylogenetic uncertainty on parameter estimation, performing the same analysis as above using a posterior distribution sample of 805 trees.

We examined how the niche trait space was occupied by Caribbean *Anolis* lizards during its diversification using disparity through time (DTT) plots (Harmon *et al.*, 2003). DTT plots estimate the average niche disparity among subclades relative to the total disparity in the entire phylogeny and we compared this observed trajectory against a mean expectation from a null model generated under Brownian motion. The disparity was calculated as the average pairwise Euclidean distance for each niche trait for the entire tree and then for subclades defined by nodes on that tree. Values approaching 0 indicate that niche variation is partitioned as among-subclade differences, suggesting that subclades exhibit little niche variation in comparison with the entire clade. Conversely, values approaching 1 indicate that subclades contain substantial niche variation and can exhibit a high niche overlap occupying similar regions in climatic space (i.e., niche convergence). Finally, it computes a niche disparity index (NDI; MDI in Harmon *et al.*, 2003) as the overall difference in the relative disparity of a clade compared to a null hypothesis simulated with a Brownian motion model. Negative NDI values indicate disparity is less than expected under a Brownian motion null model. These negative values have been interpreted as disparity accumulated early in the history of a clade (i.e., early-burst trait model). In contrast, positive NDI values indicate disparity is greater than expected under a null model, suggesting that subclades overlap more in their occupation of ecological niche space (Harmon *et al.*, 2003). Here, we generated DTT plots for species’ niche mean position and calculated the NDI index using the MCC-tree. We compared the NDI index against a mean expectation of 1000 simulated values under a BM model.

### Ecological niche modeling

We generated ecological niche models across the entire Caribbean region for 72 species. First, we estimated potential geographical distributions for each species across the Greater and Lesser Antilles islands using three algorithms (GAM: Generalized Additive Model; SRE: Surface Range Envelope; and MaxEnt: Maximum Entropy) implemented in the Biomod2 library in R (Thuiller *et al.*, 2009). We used all 19 bioclimatic variables from the WorldClim database with a resolution of 30 arc-seconds (Hijmans *et al.*, 2005) and default model parameters. Validation metrics for each model showed that Maxent models had the higher scores and we decided to use this algorithm for the following procedures (Figure S1). We generated potential geographical distributions in Maxent models using 50 pseudo-replicates by splitting the dataset into training (75%) and testing (25%) sets of occurrence points. Binary predictions (i.e., presence-absence) were generated using the 10^th^ percentile threshold as a cut-off (Liu *et al.*, 2005). Pixels predicted as present were classified as native (occupied sites) or target (non-occupied sites) given the current native distribution for each *Anolis* species modeled (Losos, 2009).

### Climatic niche structure in single-island endemics

For each potential geographical distribution, we characterized its environmental suitability using climate information from all 19 variables from Worldclim (Hijmans *et al.*, 2005). We extracted climatic information for each pixel defined as a presence from the binary predictions and then each variable was standardized. We calculated the Euclidean distance from each pixel to the niche centroid, which represents the most suitable conditions for a species, and suitability decreases as distances to the niche centroid increase (Maguire, 1973; Martínez-Meyer *et al.*, 2013; Lira-Noriega & Manthey, 2014). If climatic niche conservatism played a role in the distribution of single-island endemic species, we expected to find shorter Euclidean distances within native areas than target areas. We compared Euclidean distances between source islands versus target islands controlling for the potential confounding factors of island area and distance between islands. Also, we compared suitability values from continuous Maxent outputs between native and target islands controlling by area and distance.

We estimated climate similarity (E-space) between the Caribbean islands with the aim to establish how much of its climatic conditions were unique or shared between islands. If each island exhibits unique climate conditions, this likely have limited the potential successful colonization of other invading species. The similarity between native and target islands was evaluated with the multivariate environmental similarity surface metric (MESS; Elith *et al.*, 2010) for each species across the Caribbean region. Negative values in MESS represent nonanalog climate conditions (i.e., novel climate conditions) and positive values represent analog conditions (i.e., similar climate conditions). We expected to find negative values outside native areas as a proxy of unsuitable conditions for successful population establishment.

### Phylogenetic signal in potential colonization ability

We classified species for which we modeled their ecological niches according to their potential successful colonization ability given the predicted climate conditions outside their native areas. Whether species exhibit similar or large Euclidean distances (calculated through a one-tailed t-test) between target and native islands, we coded these as potential to population establishment regardless the ocean barriers to dispersal. In contrast, whether species exhibit small Euclidean distances or not predicted any pixels outside their native areas, we coded these as restricted by climate to its native areas. We used this new binary trait (colonizing ability vs. non-colonization ability) and measure its phylogenetic signal using the D-statistic (Fritz & Purvis, 2010), a measure of phylogenetic signal in a binary trait, with the *caper* R package (Orme *et al.*, 2013).

## Results

### Climatic niche convergence and niche disparity through time

The best final model included 29 regime shifts (*k* = 29), 11 distinct regimes (*k’* = 11) and 27 convergent shifts (c = 27) occupied by multiple lineages (Figure 1; Table 1). We discovered nine (9) convergent climate regimes (colored large dots) and two (2) non-convergent regimes (black or gray large dots; Figure 1). Most niche shifts discovered by SURFACE suggest that incursions in climate space have occurred along the precipitation axis. Only three lineages evolved by convergence to occupy cold climate regimes (red dots, Figure 1). It is likely that many niche shifts are associated with dispersal across islands. For instance, the *roquet* series of *Anolis* lizards (*A. luciae*, *A. bonairensis*, *A. griseus*, *A. trinitatis*, *A. richardi*, *A. aeneus*, *A. extremus*, and *A. roquet*) evolved to occupy a wet and hot climatic regime (light blue points, Figure 1). Similarly, four species independently evolved to occupy very rainy conditions in the climate space (*A. occultus*, *A. cuvieri*, *A.gundlachi*, and *A. evermanni*; light green dots, Figure 1). Finally, at least four lineages have evolved to occupy very hot and dry conditions (*A. bonairensis*, *A. smallwoodi*, *A. strahmi*, *A. longitibialis*, and *A. ravitergum*; ocher color dots, Figure 1). The resulting pattern of climatic niche convergence is different from an expected by a Brownian motion process (*p*-value < 0.01). A similar pattern of niche shifts across were identified using the *l*lou method (Khabbazian *et al.*, 2016; Table S1, Figure S2).

**Figure 1.**
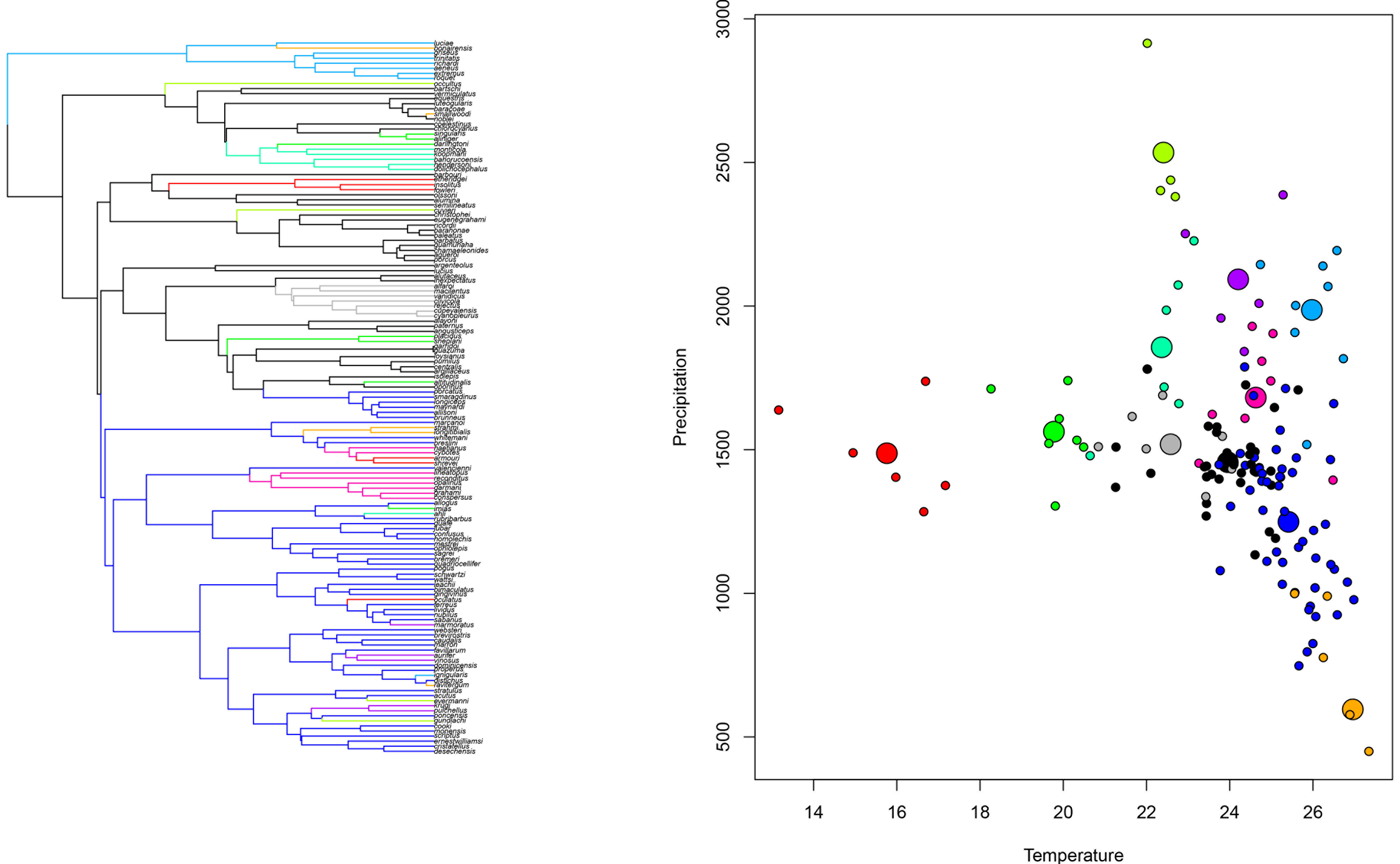
Convergent evolution of climatic niche in the Caribbean *Anolis* lizards. a) Calibrated phylogeny for Caribbean *Anolis* lizards with climatic niche data and adaptive regimes identified using a SURFACE approach. Colored branches represent convergent adaptive peaks (or climate regimes) and gray or black branches represent non-convergent regimes. b) Climatic space occupied by the Caribbean *Anolis* species. Large colored dots represent the same convergent adaptive regimes identified in the phylogeny and small colored dots represent the species that have evolved around these adaptive optima. Large gray or black dots represent the same non-convergent regimes identified in the phylogeny. Small gray or black dots correspond to species that have evolved toward these regimes.

**Table 1.** Convergence parameters from each regime identified by SURFACE in the backward phase (19 regimes) based on the Akaike Information Criteria. Δ AICc: corrected Akaike Information Criterion; Δ AICc: corrected Delta Akaike Information Criterion; k’: Number of distinct climate regimes after collapsing convergent regimes; Δk: Reduced number of regimes after accounting for climatic niche convergence; c: number of shifts that are towards convergent regimes occupied by multiple lineages; k’ conv: Number of convergent regimes reached by multiple shifts. Selected best model in bold.

| Regime | AICc | $\Delta$ AICc | k' | Δk | c | k' conv |
| --- | --- | --- | --- | --- | --- | --- |
| 1 | 2502 | 104 | 29 | 0 | 0 | 0 |
| 2 | 2493 | 95 | 28 | 1 | 2 | 1 |
| 3 | 2485 | 87 | 27 | 2 | 4 | 2 |
| 4 | 2477 | 79 | 26 | 3 | 6 | 3 |
| 5 | 2469 | 71 | 25 | 4 | 7 | 3 |
| 6 | 2461 | 63 | 24 | 5 | 9 | 4 |
| 7 | 2453 | 55 | 23 | 6 | 11 | 5 |
| 8 | 2446 | 48 | 22 | 7 | 13 | 6 |
| 9 | 2439 | 41 | 21 | 8 | 15 | 7 |
| 10 | 2432 | 34 | 20 | 9 | 16 | 7 |
| 11 | 2426 | 28 | 19 | 10 | 17 | 7 |
| 12 | 2421 | 23 | 18 | 11 | 19 | 8 |
| 13 | 2416 | 18 | 17 | 12 | 21 | 9 |
| 14 | 2411 | 13 | 16 | 13 | 22 | 9 |
| 15 | 2407 | 9 | 15 | 14 | 24 | 10 |
| 16 | 2402 | 4 | 14 | 15 | 24 | 9 |
| 17 | 2400 | 2 | 13 | 16 | 25 | 9 |
| 18 | 2398 | 1 | 12 | 17 | 26 | 9 |
| **19** | **2398** | **0** | **11** | **18** | **27** | **9** |

The rate of adaptation to optima (α) for temperature was 0.660 and for precipitation was 0.628. Conversely, the rate of stochastic evolution (σ^2^) was higher for precipitation (5035.9) than for temperature (1.059). The expected time to evolve halfway to an optimum, calculated in millions of years (*t*_1/2_ or log(2)/ α), was similar for both climate axes (temperature = 1.05 and precipitation = 1.10).

The plots of disparity over time for temperature and precipitation exhibited different trajectories through the Caribbean *Anolis* radiation. Niche disparity for temperature was different than expected under a Brownian motion model (left panel in Figure 2). Niche disparity for precipitation exhibited a trajectory different than expected under a Brownian motion model, but the disparity values were highly positives (right panel in Figure 2). For the for temperature and precipitation the niche disparity values calculated for all the tree were positive (temperature NDI=0.151, p<0.00; precipitation NDI=0.471; p<0.001).

**Figure 2.**
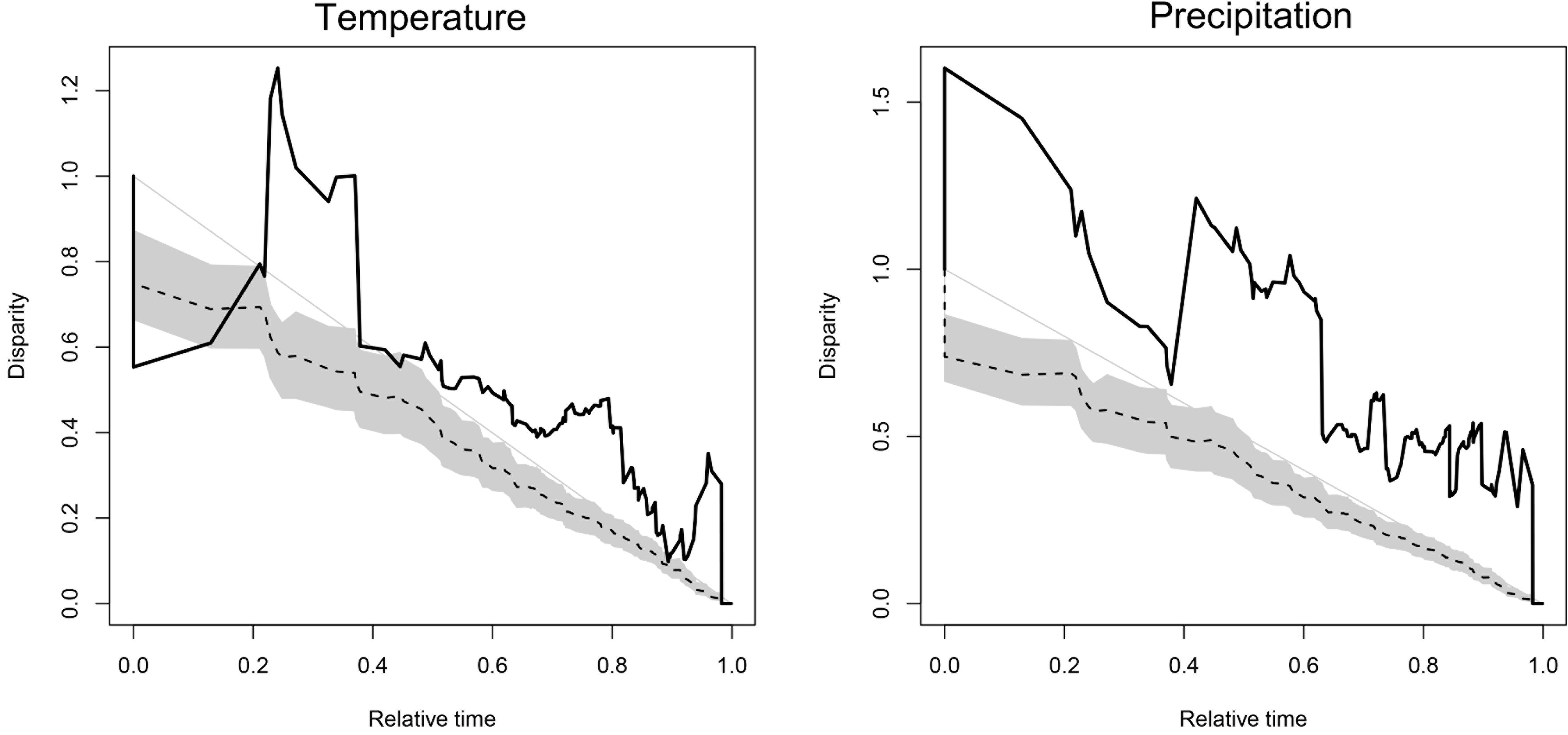
Disparity through time plots (DTT) for temperature and precipitation for the Caribbean *Anolis* lizards. Relative time indicates the entire time span of the phylogeny from 0 (i.e., root) to 1 (i.e., tips). The dashed line indicates the median DTT based on 1000 simulations of each trait under a Brownian motion model. The gray shaded area indicates the 95% DTT range for the simulated data.

### Climatic niche structure in single-island endemics

Target islands (non-occupied islands) exhibited higher values in Euclidean distances to niche centroid than native islands in single-islands (*p*-value < 0.001; Figure 3). From 73 species used in these analyses, only 14 species showed that target and native islands did not differ in their Euclidean distances to niche centroid (Table S2). Based on this result, we coded species as with potential to colonize other islands and species limited by available climatic conditions. The phylogenetic signal of this binary trait was low (D-statistic = 0.36; *p*-value = 0.014). However, this D-statistic value was not different from an expected by a null Brownian motion model (*p*-value=0.22).

**Figure 3.**
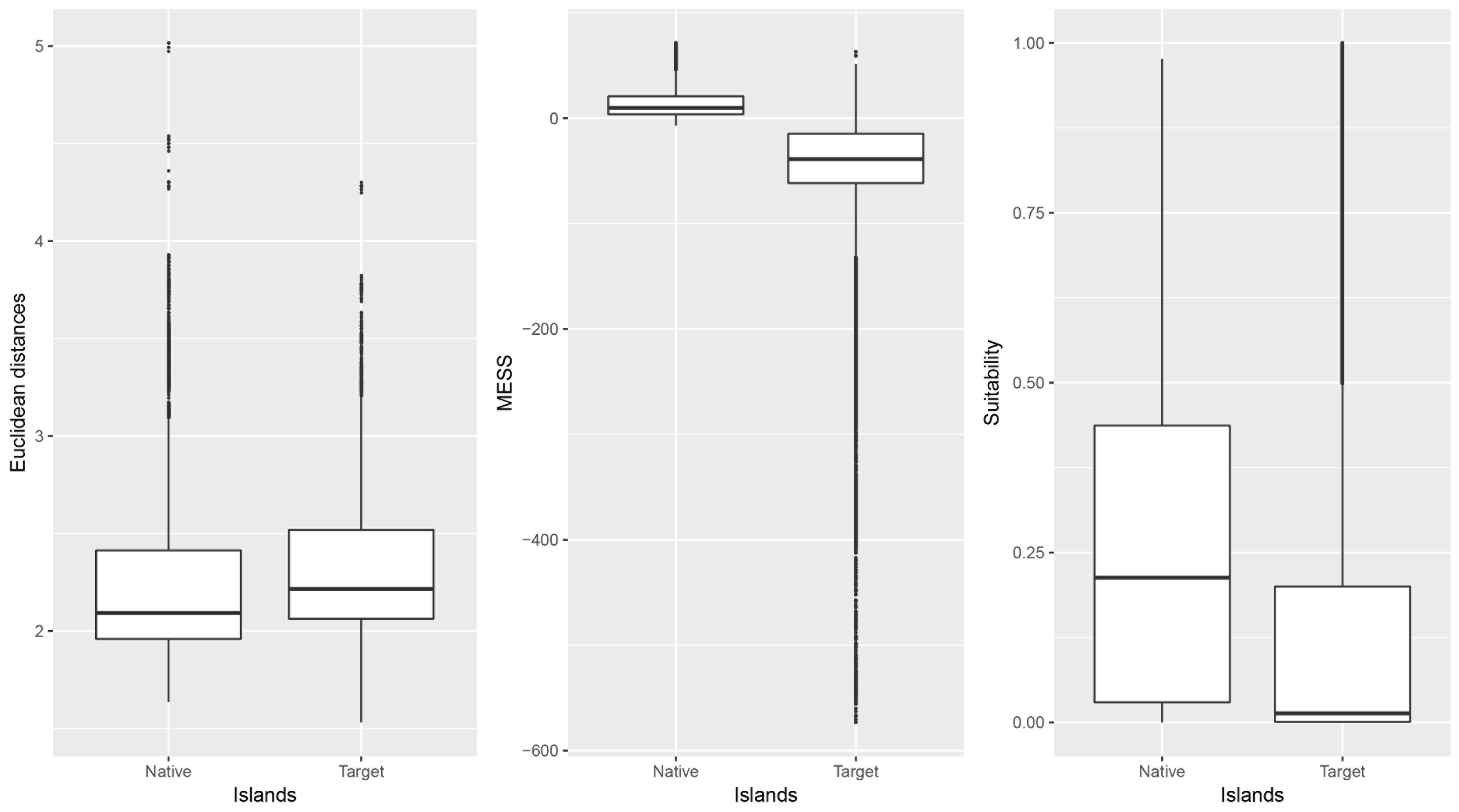
Boxplots of climatic niche structure (Ecological distances), multivariate environmental similarity surface (MESS) and climatic suitability (Maxent’s logistic outputs) between native (occupied) islands and target (non-occupied) islands. See main text for details.

MESS analyses showed that novel climate conditions (i.e., negative values) are spatially concentrated outside native islands for single-endemic anole lizards (*p*-value < 0.001; Figure 3). Finally, suitability values from Maxent models showed that single-island endemics tend to exhibit low values outside their native areas (*p*-value < 0.001; Figure 3).

## Discussion

Climatic niche convergence and niche conservatism have been two very pervasive patterns throughout the Caribbean *Anolis* radiation. Our results show that climatic niche disparity (i.e., niche divergence) has decreased over time. Also, we detected several instances of niche convergence through time (i.e., dispersed through the entire anole phylogeny) and across space (i.e., among islands). These niche convergence events likely occurred as species dispersed or were isolated at the major Caribbean landmasses (Losos, 2009).

Both climatic niche stasis and shifts apparently have been recurrent across the entire Caribbean anole radiation. For instance, the *Ctenonotus* clade distributed in Lesser Antilles (*A. bimaculatus* group) has retained its ancestral niche occupying a hot and dry climate regime (Figure 1). However, some species in this group (e.g., *A. oculatus*) evolved to occupy cooler and humid conditions than its most recent common ancestor. Some niche shifts are likely associated to dispersal events between islands or within islands.

Recent divergence time estimates for *Anolis* lizards suggest that anoles colonized the Caribbean islands (or GAARlandia; Poe *et al.*, 2017) before the Eocene-Oligocene glacial maximum. During that time, the global climate was warmer and drier than it is today (i.e., the Middle Miocene Climatic Optimum; Zachos *et al.*, 2008). Anoles likely evolved toward cooler and wetter climate conditions after they colonized the Caribbean landmasses and as clades isolated due to the break-up of Caribbean landmass likely occupied available climate spaces quickly in each island (Velasco *et al.*, 2016). Parameter estimates of SURFACE showed that the rate of evolution in the thermal niche dimension was slower than hydric one. This suggests that anoles have been more labile to occupy a precipitation gradient than a temperature gradient in the Caribbean islands. Disparity trends suggest that niche evolution has been pervasive across the entire radiation showing higher values than expected under a BM model. Rapid niche evolution can be facilitated by dispersal between and within islands. A previous study found that rates of climatic niche evolution vary with the geographical area and climate heterogeneity (Algar & Mahler, 2016)

The observation that several clades of Caribbean anoles diversified extensively in isolation and converged to occupy similar niches suggests that this might be the result of adaptation towards similar environmental regimes (Boucher *et al.*, 2011). It is likely that strong stabilizing selection would operate in climatic niches in these single-island endemics (Holt & Gaines, 1992; Holt, 1996). This is supported by our finding that optimal environmental requirements for single-island endemics were low and scarce outside their native islands. This supports the hypothesis that climatic niche adaptation has been pervasive across Caribbean anole radiation and has played an important role in shaping the exceptional insular endemism observed today: only 16 out of 166 species have colonized more than one island (Helmus *et al.*, 2014). Finally, the observed niche shifts towards cooler temperatures in the Caribbean region seem to represent an independent evolutionary adaptation. These independent adaptations in some lineages toward cooler conditions have been faster, as in the *cybotes* clade in Hispaniola (Muñoz *et al.*, 2014).

Regarding the recent criticism that Akaike information criterion (AIC) tends to inflate the false rate of discovery of convergent regimes in comparative datasets (Khabbazian *et al.*, 2016), we consider that the alternative procedure presented by Khabbazian *et al.*, (2016) based on a Bayesian information criterion (BIC) tends to penalize in excess model complexity (underfit) and therefore always selects very conservative convergent regimes. Because AIC and BIC are based on different philosophies and mathematical formulation, it is not surprising that they select different models. As our aim here is discovery (i.e., prediction), we consider that AIC is a good criterion for this purpose. Regardless, using AIC in SURFACE and *l*lou, we recovered very similar convergent climatic regimes in Caribbean *Anolis* lizards.

Finally, we acknowledge that our niche estimates are based on occurrence data and therefore are only an approximation to the fundamental niche (Hutchinson, 1957; Soberón, 2007). Although some researchers argue that estimates of physiological tolerances (e.g., critical thermal maximum and minimum) would be a better approach to evaluate niche dimensions; however, these parameters do not capture the true fundamental niche either, because this can only be estimated using *in situ* population growth rate information (Pulliam, 2000; Holt, 2009). Accordingly, it would be necessary to establish whether occurrence data or physiological tolerance limits more closely estimate the Hutchinsonian fundamental niche.

### Caribbean anole endemicity

Recent work on anole biogeography have shown that dispersal has prevailed in the evolutionary history of Caribbean anoles (Poe *et al.*, 2017). Most inferred past dispersal events have occurred when Caribbean islands were merged in a single landmass (Poe *et al.*, 2017). Conversely, recent dispersal events across the ocean have been facilitated by humans (Powell *et al.*, 2011; Helmus *et al.*, 2014). In these recent cases, impoverished islands in anole diversity have been easily colonized by different anole species (Helmus *et al.*, 2014).

These results suggest that dispersal ability might be prevalent in insular anole lizards, but ocean barriers to dispersal have limited the successful colonization of more Caribbean islands. Our results might contradict this idea and suggest that not all species have the capacity to colonize other islands regardless the geographic barriers. Our results suggest that environmental suitability tend to be limited outside native islands for many single-island endemic *Anolis* species. In addition, MESS analysis also suggests that each Greater Antillean island exhibits a unique set of climatic conditions (Williams & Jackson, 2007; Velasco *et al.*, 2016) which would limit the establishment of potential colonizers. The D-statistic showed that phylogenetic signal in the potential ability to colonize given the availability of suitable climatic conditions was low and suggests a phylogenetic clumping (Fritz & Purvis, 2010). This implies that climatic suitability beyond native areas can be predicted by phylogenetic affinity as it has been found with morphological traits (Latella *et al.*, 2011; Poe *et al.*, 2011)

Although single-island endemics might have truncated climatic niches (i.e., species with larger niches than observed but truncated by geography), their niche requirements are also not available in the surrounding geography, as the climatic similarity metrics revealed (Figure 3). However, our results suggest that small Caribbean islands or banks might be good candidates for the establishment of colonizing species because they tend to hold higher values of climatic suitability (Petitpierre *et al.*, 2012). All these results suggest that climatic suitability has played an important role, coupled with ocean barriers, in limiting the successful colonization of species to other islands. Many invasion events occurring in small islands (Helmus *et al.*, 2014, 2016) and a few in large islands (e.g., *A. sagrei* in Jamaica) support this conclusion. However, we recommend that other approaches, either based on physiological tolerances or demographic data (Kearney & Porter, 2009; Kolbe *et al.*, 2014) should be implemented to corroborate or not our results about novel climates limiting dispersal and successful population establishment in the Greater Antilles.

### Conclusions

Our results suggest that climatic niche divergence occurred early in the Caribbean *Anolis* radiation involving multiple episodes of niche shifts, particularly six large events of climatic niche convergence. Caribbean anoles have evolved repeatedly across the climate space occupying different combinations of temperature and precipitation. Multiple lines of evidence suggest that niche divergence has been pervasive along time and across space, but more recent niche conservatism is likely to have played a role driving insular endemism in *Anolis* lizards due to climatic dissimilarity across the Caribbean islands. A next step requires corroborating these main finding using other experimental or modeling approaches and test whether climate dissimilarity also has operated as a driver of insular endemism in other taxonomic groups. We consider that a combination of approaches provides stronger lines of evidence for testing hypotheses about the causes of insular endemism.

## Acknowledgments

This study was made possible thanks to the devoted effort of all Caribbean herpetologists, particularly those working with anoles, who have documented the diversity of Caribbean *Anolis* lizards over the last 300 years. The first author thanks the Posgrado de Ciencias Biológicas (PCB) program at the Universidad Nacional Autónoma de Mexico (UNAM), and the Consejo Nacional de Ciencia y Tecnología (Conacyt) for a graduate scholarship. The Faculty of Sciences at UNAM and a DGAPA postdoctoral grant facilitated the culmination of this manuscript.

## References

Algar, A.C. & Mahler, D.L. 2015. Area, climate heterogeneity, and the response of climate niches to ecological opportunity in island radiations of Anolis lizards. Glob. Ecol. Biogeogr. 25, 781–791.

Bacon, C.D., Baker, W.J. & Simmons, M.P. 2012. Miocene dispersal drives island radiations in the palm tribe Trachycarpeae (Arecaceae). Syst Biol, 61, 426–42.

Boucher, F.C., Thuiller, W., Roquet, C., Douzet, R., Aubert, S., Alvarez, N. & Lavergne, S. 2011. Reconstructing the origins of high-alpine niches and cushion life form in the genus Androsace s.l. Primulaceae. Evolution, 66, 1255–1268.

Bravo, G.A., Remsen Jr., J. V. & Brumfield, R.T. 2014. Adaptive processes drive ecomorphological convergent evolution in antwrens Thamnophilidae. Evolution, 68, 2757–2774.

Brown, J.H. & Lomolino, M. V. 1998. Biogeography, 2nd ed. Sinauer Associates, Inc. Sunderland, Massachusetts.

Butler, M.A. & King, A.A. 2004. Phylogenetic comparative analysis: a modeling approach for adaptive evolution. Am. Nat. 164, 683–695.

Donoghue, M.J & Edwards, E. J. 2014. Biome shifts and niche evolution in plants. Annu. Rev. Ecol. Evol. Syst. 45, 547–572.

Elith J., Kearney M., & Phillips S. 2010. The art of modelling range-shifting species. Methods Ecol. Evol. 1, 330–342.

Fritz, S.A., & Purvis, A. 2010. Selectivity in mammalian extinction risk and threat types: anew measure of phylogenetic signal strength in binary traits. Conserv. Biol. 24, 1042–1051.

Harmon, L.J., Schulte, J. A, Larson, A. & Losos, J.B. 2003. Tempo and mode of evolutionary radiation in iguanian lizards. Science, 301, 961–4.

Helmus,M.R., Mahler, D.L. & Losos, J.B. 2014. Island biogeography of the Anthropocene. Nature, 513, 543–546.

Helmus, M. R., Behm, J. E., Jesse, W. A., Kolbe, J. J., Ellers, J., & Losos, J. B. 2016. Exotics exhibit more evolutionary history than natives. In: Invasion Genetics: The Baker and Stebbins Legacy (eds S. C. H. Barrett, S.C.H., Colautti, R.I., Dlugosch, K.M & Rieseberg, L.H., eds), pp. 122–138. John Wiley & Sons, Ltd, Chichester, UK.

Hijmans, R.J., Cameron, S.E., Parra, J.L., Jones, P.G. & Jarvis, A. 2005. Very high resolution interpolated climate surfaces for global land areas. Int. J. Climatol. 25, 1965–1978.

Holt, R.D. 1996. Demographic constraints in evolution: towards unifying the evolutionary theories of senescence and niche conservatism. Evol. Ecol. 10, 1–11.

Holt, R. D. 2009. Bringing the Hutchinsonian niche into the 21st century: ecological and evolutionary perspectives. PNAS 106, 19659–19665.

Holt, R.D. & Gaines, M.S. 1992. The analysis of adaptation in heterogeneous landscapes: implications for the evolution of fundamental niches. Evol. Ecol. 6, 433–447.

Hosner, P.A., Sánchez-González, L.A., Peterson, A.T., & Moyle,R.G. 2014. Climate-driven diversification and Pleistocene refugia in Philippine birds: evidence from phylogeographic structure and paleo-environmental niche modeling. Evolution, 68, 2658–2674.

Hutchinson, G.E. 1957. Concluding remarks. Cold Spring Harb. Symp. Quant. Biology. 22, 415–427.

Ingram, T. & Mahler, D.L. 2013. SURFACE: detecting convergent evolution from comparative data by fitting Ornstein-Uhlenbeck models with stepwise Akaike Information Criterion. Methods Ecol. Evol. 4, 416–425.

Kearney, M. & Porter, W. 2009. Mechanistic niche modelling: combining physiological and spatial data to predict species’ ranges. Ecol. Lett. 12, 334–350.

Khabbazian, M., Kriebel, R., Rohe, K., & Ané, C. 2016. Fast and accurate detection ofevolutionary shifts in Ornstein-Uhlenbeck models. Methods Ecol. Evol. 7, 811–824.

Kier, G., Kreft, H., Lee, T.M., Jetz, W., Ibisch, P.L., Nowicki, C., Mutke, J. &Barthlott, W. 2009. A global assessment of endemism and species richness across island and mainland regions. PNAS 106, 9322–7.

Kolbe, J. J., Ehrenberger, J. C., Moniz, H. A. & Angilletta Jr, M. J. 2013. Physiological variation among invasive populations of the brown anole (Anolis sagrei). Physiol. Biochem. Zool. 87, 92–104.

Latella, I. M., Poe, S., & Giermakowski, J. T. 2011. Traits associated with naturalization in Anolis lizards: comparison of morphological, distributional, anthropogenic, and phylogenetic models. Biol. Invasions 13, 845–856.

Lira- Noriega, A. & Manthey J.D. 2014. Relationship of genetic diversity and niche centrality: a survey and analysis. Evolution 68: 1082–1093.

Liu, C., Berry, P.M., Dawson, T.P. & Pearson, R.G. 2005. Selecting thresholds of occurrence in the prediction of species distributions. Ecography, 3, 385–393.

Losos, J.B. 2009. Lizards in an Evolutionary Tree: Ecology and Adaptive Radiation of Anoles, University of California Press, Berkeley, CA.

Losos, J.B. & Ricklefs, R.E. 2010. The theory of island biogeography revisited, Princeton University Press, Princeton and Oxford.

MacArthur, R.H. & Wilson, E.O. 1967. The theory of island biogeography, Princeton University Press. Princeton.

Mahler, D.L., Ingram, T., Revell, L.J. & Losos, J.B. 2013. Exceptional convergence on the macroevolutionary landscape in island lizard radiations. Science, 341, 292–5.

Martínez-Meyer, E., Díaz-Porras, D., Peterson, A.T. & Yáñez-Arenas, C. 2013. Ecological niche structure and rangewide abundance patterns of species. Biol. Lett. 9, 20120637.

Maguire Jr. B.,1973. Niche response structure and the analytical potentials of its relationship to the habitat. Am. Nat. 213–246.

Muñoz, M.M., Stimola, M.A., Algar, A.C., Conover, A., Rodriguez, A.J., Landestoy, M.A., Bakken, G.S. & Losos, J.B. 2014. Evolutionary stasis and lability in thermal physiology in a group of tropical lizards. Proc. R. Soc. Lond. 281, 20132433.

Orme, D., Freckleton, R., Thomas, G., Petzoldt, T., Fritz, S., Isaac, N & Pearse, W. 2013. caper: Comparative Analyses of Phylogenetics and Evolution in R. R package version 0.5.2. https://CRAN.R-project.org/package=caper

Peterson, A.T., Soberon, J., Pearson, R.G.,Anderson, R.P., Martinez-Meyer, E., Nakamura,M. & Araujo, M.B. 2011. Ecological Niches and Geographic Distributions, Princeton University Press.

Petitpierre, B., Kueffer, C., Broennimann, O., Randin, C., Daehler, C., & Guisan, A. 2012. Climatic niche shifts are rare among terrestrial plant invaders. Science, 335, 1344–1348.

Poe, S., Giermakowski, J. T., Latella, I., Schaad, E. W., Hulebak, E. P., & Ryan, M. J. 2011. Ancient colonization predicts recent naturalization in Anolis lizards. Evolution 65, 1195–1202.

Poe, S., Nieto-Montes de Oca, A., Torres-Carvajal,O., de Queiroz, K., Velasco, J.A., Truett, Gray, L.,B., Ryan M.J., Kohler, G., Ayala-Varela & Latella, I. A phylogenetic, biogeographic, and taxonomic study of all extant species of Anolis (Squamata; Iguanidae). Syst. Biol. https://doi.org/10.1093/sysbio/syx029

Powell, R., Henderson, R.W., Farmer, M.C., Breuil, M., Echternacht, A.C., van Buurt, G., Romagosa, C.M. & Perry, G. 2011. Introduced amphibians and reptiles in the Greater Caribbean: Patterns and conservation implications, pp. 63–143. In Conservation of Caribbean Island Herpetofaunas. Volume 1 (Hailey, A., Wilson, B.S. & Horrocks, J.A. eds). Brill, Leiden, The Netherlands.

Pulliam, H.R. 2000. On the relationship between niche and distribution. Ecol. Lett. 3, 349–361.

Qian, H. & Ricklefs, R.E. 2012. Disentangling the effects of geographic distance and environmental dissimilarity on global patterns of species turnover. Glob. Ecol. Biogeogr., 21, 341–351.

Revell, L.J., Harmon, L.J. & Collar, D.C. 2008. Phylogenetic signal, evolutionary process, and rate. Syst. Biol., 57, 591–601.

Soberón, J. 2007. Grinnellian and Eltonian niches and geographic distributions of species. Ecol. Lett. 10, 1115–1123.

Stuart, Y.E., Losos, J.B. & Algar, A.C. 2012. The island-mainland species turnover relationship. Proc. R. Soc. Lond., 279, 4071–4077.

Thuiller, W., Lafourcade, B., Engler, R. & Araújo, M.B. 2009. BIOMOD - a platform for ensemble forecasting of species distributions. Ecography, 32, 369–373.

Velasco, J.A., Martínez-Meyer, E., Flores-Villela, O., García-Aguayo, A., Algar, A.C. & Kohler, G. 2016. Climatic niche attributes and diversification in Anolis lizards. J. Biogeogr. 43: 134–144.

Wiens, J. J., & Graham, C.H. 2005. Niche conservatism: integrating evolution, ecology, and conservation biology. Annu. Rev. Ecol. Evol. Syst. 36, 519–539.

Williams, J.W. & Jackson, S.T. 2007. Novel climates, non-analog communities, and ecological surprises. Front. Ecol. Environ. 57, 475–482.

Wüest,R.O., Antonelli, A., Zimmermann, N.E. & Linder, H.P. 2015. Available climate regimes drive niche diversification during range expansion. Am. Nat. 5, 640–652.

Zachos, J.C., Dickens, G.R. & Zeebe, R.E. 2008. An early Cenozoic perspective on greenhouse warming and carbon-cycle dynamics. Nature, 451, 279–83.

